## Supplementary material for "Inhibition of mitochondrial dynamics preferentially targets pancreatic cancer cells with enhanced tumorigenic and invasive potential": Figure S1, S2, S3, S4

### Figure S1, related to Figure 1

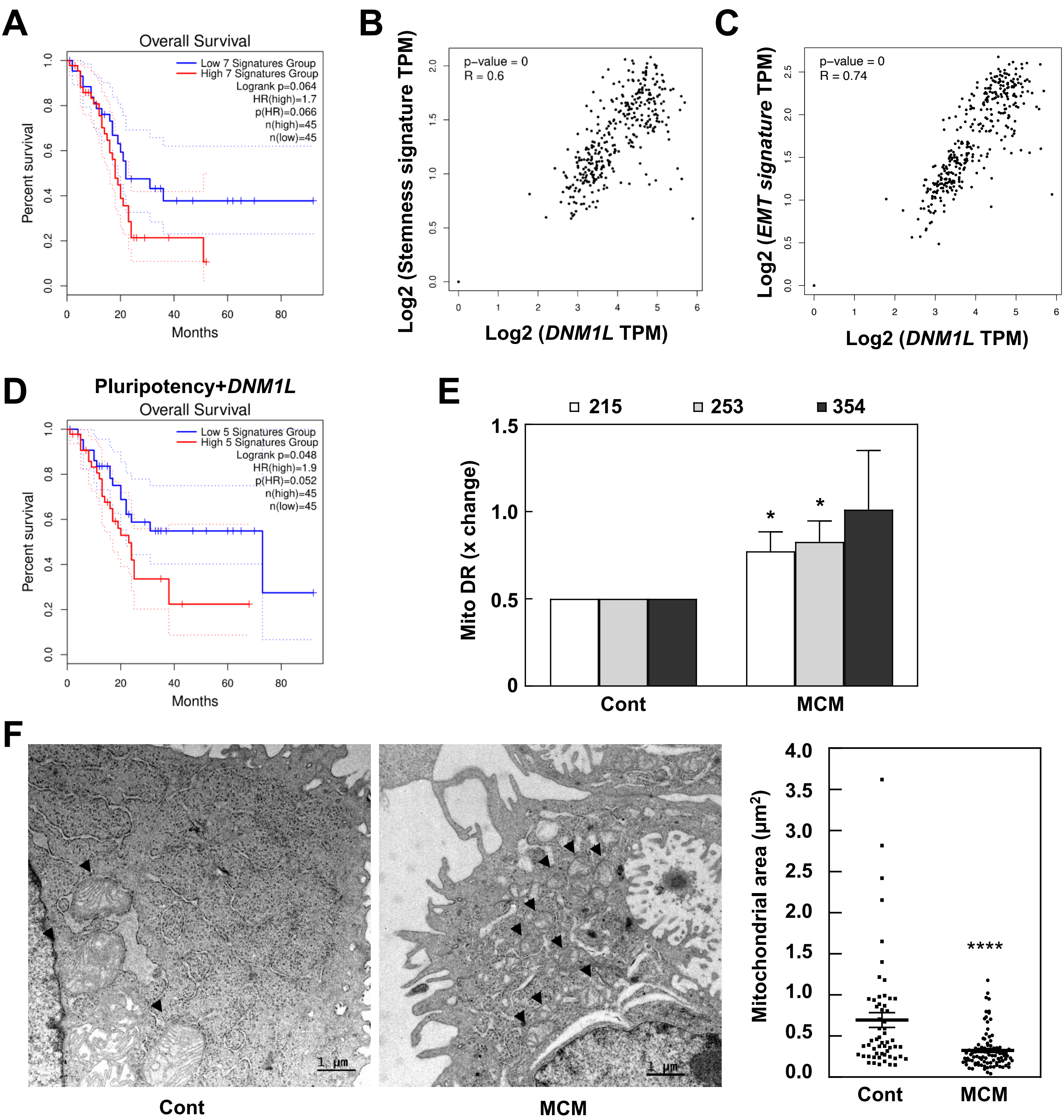


**Figure S1. Mitochondrial fission relates to stemness and EMT in human PDAC. A.** Overall survival of PDAC patients in the upper and lower quartiles for a mitochondrial dynamics signature (*DNM1L*, *DNM2*, *FIS1*, *MFF*, *MFN1*, *MFN2* and *OPA1*). **B**, **C**. Correlation of *DNM1L* expression and stemness (*NANOG*, *OCT4*, *KLF4*, *SOX2*) (B) or EMT signatures (*ZEB1*, *SNAI1* and *SNAI2*) (C). **D.** Overall survival of PDAC patients in the upper or lower quartiles for above stemness signature combined with *DNM1L* expression. **E.** Mitochondrial mass as determined by flow cytometry using MitoTracker™ Deep Red FM in either control cells or cells treated with conditioned media from M2-polarized macrophages (macrophage-conditioned media, MCM) (n=3-4). **F.** TEM images and quantification of the mitochondrial area of control cells *vs* cells treated with MCM (n=9-12 pictures representing 57 *vs* 94 mitochondria). *p < 0.05, ****p < 0.0001 using the Mann & Whitney test.

#### Figure S2, related to Figure 2

**A**

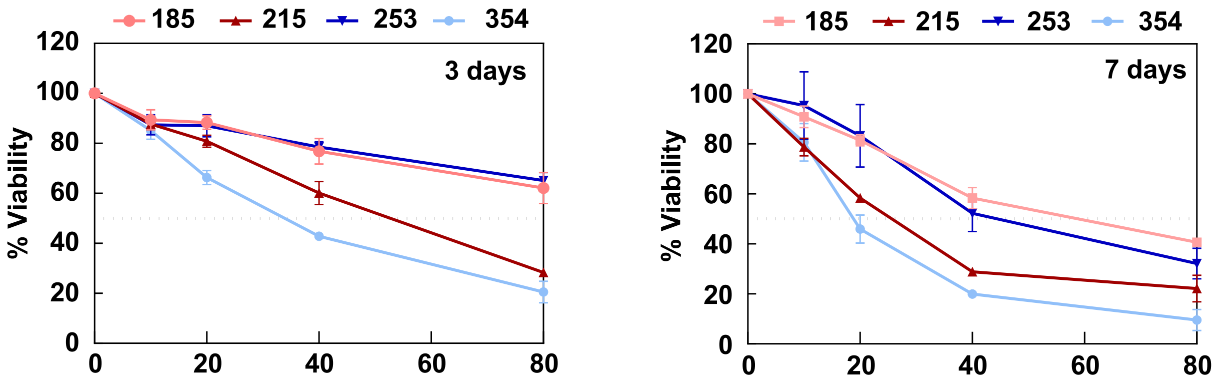


**B**

| IC50 (μM) | 3d | 7d |
|---|---|---|
| 185 | n.d. | 54.23 |
| 215 | 52.77 | 22.99 |
| 253 | n.d. | 42.43 |
| 354 | 32.51 | 18.74 |

**C**

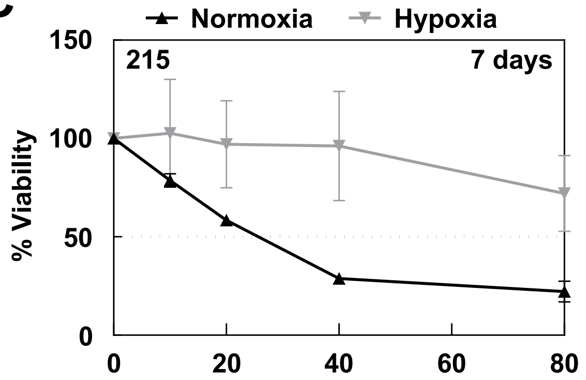


**Figure S2. mDivi-1 decreases cell viability in normoxia**. **A**, **B.** Evaluation of mDivi-1 IC50 for cell proliferation in normoxic condition for 4 different PDX models after 3 and 7 days of treatment. **C.** Evaluation of mDivi-1 IC50 for proliferation in normoxic or hypoxic conditions for 215 cells after 7 days of treatment.

### Figure S3, related to Figure 3

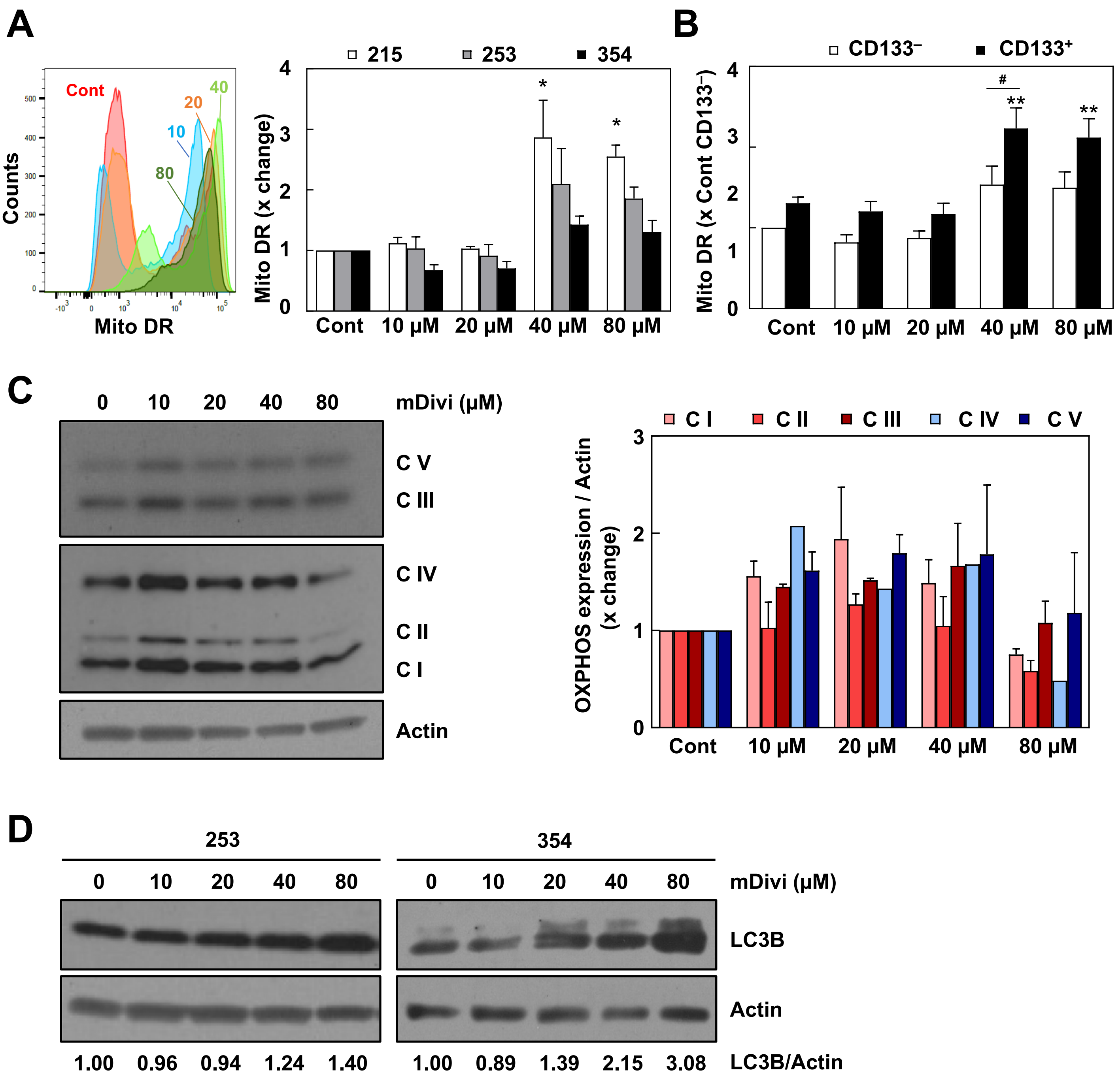


**Figure S3**. **mDivi-1 disrupts mitochondrial function. A**, **B.** Mitochondrial mass as measured by MitoTracker™ Deep Red FM staining for the bulk cell population (A) and separated for CD133⁻ *vs* CD133⁺ cells (B) (n=4-9). Panel A on the left shows a representative plot for MitoTracker™ Deep Red FM median staining for the bulk cell population. **C.** Protein expression for mitochondrial respiratory chain complexes as assessed by WB (left) and the corresponding densitometric quantification (right) for 354 cells. **D.** LC3B protein expression as assessed by WB for 253 and 354 cells. The numbers below indicate the quantification of expression relative to actin. In C, D, actin was used as loading control for densitometric analyses. * *vs* control condition; # *vs* CD133⁻ for indicated condition. *p<0.05, **p < 0.01, ***p < 0.001, ****p < 0.0001; Kruskal-Wallis with Dunn's post-test (A); ANOVA with Bonferroni *post-hoc* test (B).

### Figure S4, related to Figure 4

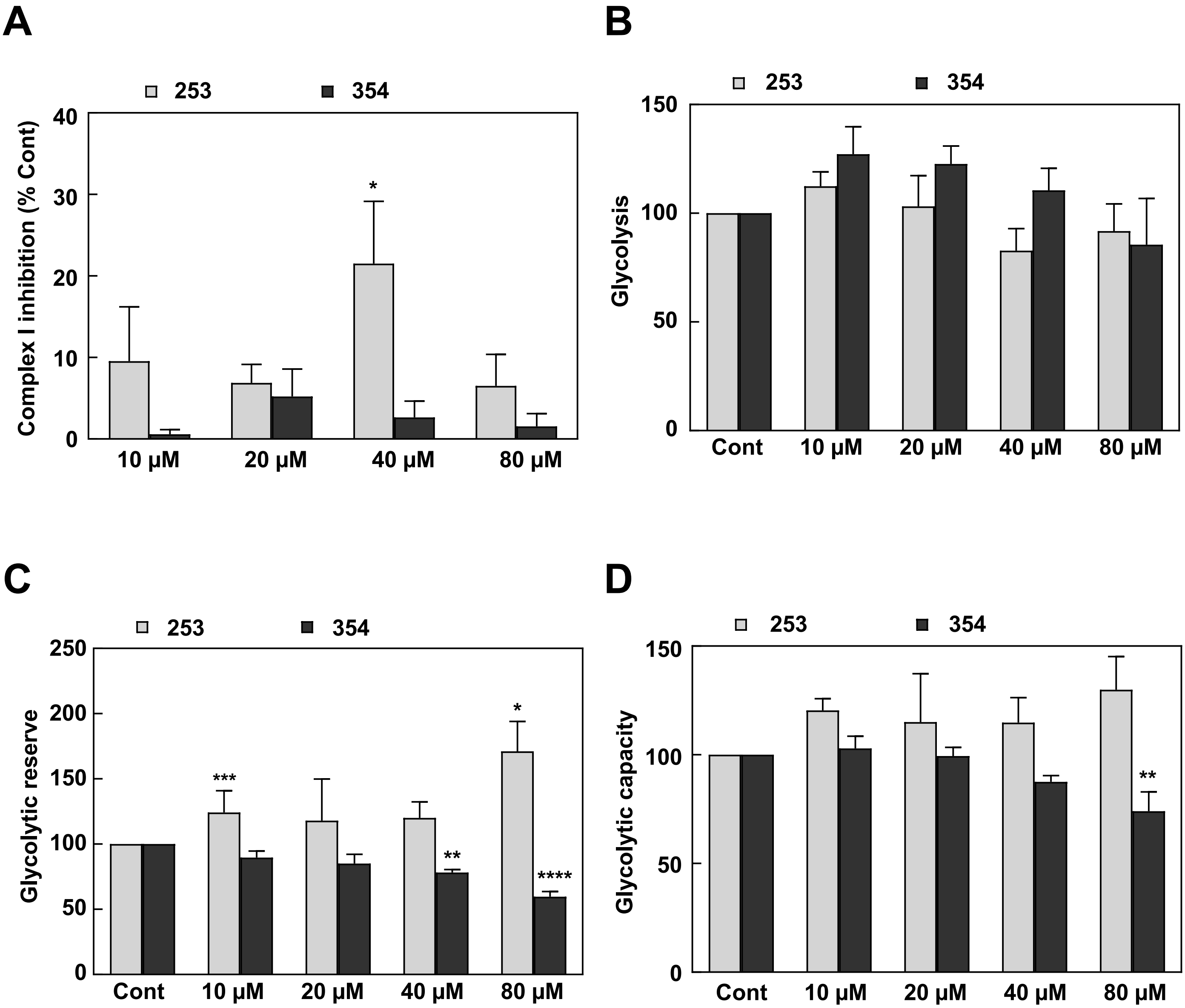


**Figure S4. Treatment effects of mDivi-1 on cellular metabolism. A.** Percentage of complex I inhibition following acute injection of mDivi-1 (n = 6). Total complex I activity was determined based on OCR inhibition by the irreversible complex I inhibitor rotenone. **B**, **C**, **D.** Measurement of glycolytic rates of the glyco stress kit in 253 and 354 cells after mDivi-1 treatment for 72h (n=3-6). *p<0.05, **p < 0.01, ***p < 0.001, ****p < 0.0001; Kruskal-Wallis with Dunn's post-test (A); ANOVA with Bonferroni *post-hoc* test (B-D).
